# Biologically Plausible Dopamine-Modulated STDP Model of Pavlovian Learning in Spiking Neural Networks

**DOI:** 10.64898/2026.07.17.739276

**Authors:** WooJun Park, Kyoung J. Lee

## Abstract

Spike-timing-dependent plasticity (STDP) and dopamine (DA) are fundamental to reward-based learning and memory formation. A widely used DA-modulated STDP model explains how neural networks associate stimuli with delayed dopaminergic rewards through an eligibility trace. However, we show that this model supports learning even at unrealistically high DA concentrations because DA simply scales the magnitude of STDP without changing its temporal profile. In contrast, experiments demonstrate that DA nonlinearly reshapes the STDP window, converting long-term depression (LTD) into long-term potentiation (LTP) at high DA levels. We therefore propose a DA-modulated STDP rule in which increasing DA progressively biases plasticity toward potentiation while receptor saturation limits further DA effects beyond a critical concentration. Simulations of recurrent networks of Izhikevich neurons show that the proposed rule supports robust conditioning only within a biologically realistic DA range (0.04–0.70 ***µ***M). Successful learning produces a hybrid network architecture consisting of a strong feedforward backbone embedded within recurrent circuitry and generates enhanced burst responses selectively to reward-associated stimuli. At the upper limit of the biologically plausible DA range, the network passes through a narrow bistable regime, converging to one of two distinct stable configurations. At higher DA concentrations, conditioning fails altogether. These results provide a biologically grounded model of DA-dependent plasticity and offer new insight into how abnormal dopamine signaling can impair learning in neurological disorders.

## 1 Introduction

DA-modulated STDP is one of the best-known implementations of the three-factor learning rule, a fundamental framework for biologically plausible synaptic plasticity. In this framework, the STDP rule modifies synaptic strength according to the relative timing of presynaptic and postsynaptic spikes, while DA acts as a neuromodulatory signal that gates synaptic plasticity (Frémaux & Gerstner, 2016). A seminal study by Izhikevich (2007) introduced a DA-modulated STDP model that associates spatially patterned stimuli with temporally delayed dopaminergic rewards in a spiking neural network. By incorporating an eligibility trace, the model enables delayed DA signals to reinforce neural activity elicited by previously presented stimuli.

Building on this framework, we recently showed that the original DA-modulated STDP rule drives the emergence of a strong feedforward organization from an initially random recurrent network (Jeong & Lee, 2025). We further demonstrated that the same learning rule can encode not only spatial patterns but also spatiotemporal sequences (Park, Kim, Jeong, & Lee, 2025), and, when the network supports superbursts, temporal intervals extending to approximately 100 ms (Kim & Lee, 2022).

Despite these successes, the original Izhikevich model has an important limitation. In the original formulation, DA acts solely as a multiplicative gain on the STDP function (Fig. 1A), uniformly scaling the magnitude of the STDP window without altering its shape or polarity (Fig. 1B). Consequently, DA regulates only the learning rate of synaptic weight updates. Experimental studies, however, have shown that DA-dependent plasticity is substantially more complex. In particular, elevated DA levels can convert long-term depression (LTD) into long-term potentiation (LTP) (Brzosko, Schultz, & Paulsen, 2015; Zhang, Lau, & Bi, 2009). Rather than simply increasing the magnitude of synaptic plasticity, elevated dopamine can alter the intracellular signaling state of the postsynaptic neuron, biasing the induction of synaptic plasticity toward LTP and, under appropriate conditions, converting spike-timing protocols that normally produce LTD into LTP. (Calabresi, Picconi, Tozzi, & Di Filippo, 2007; Pawlak & Kerr, 2008; Shen, Flajolet, Greengard, & Surmeier, 2008)

**Fig. 1.**
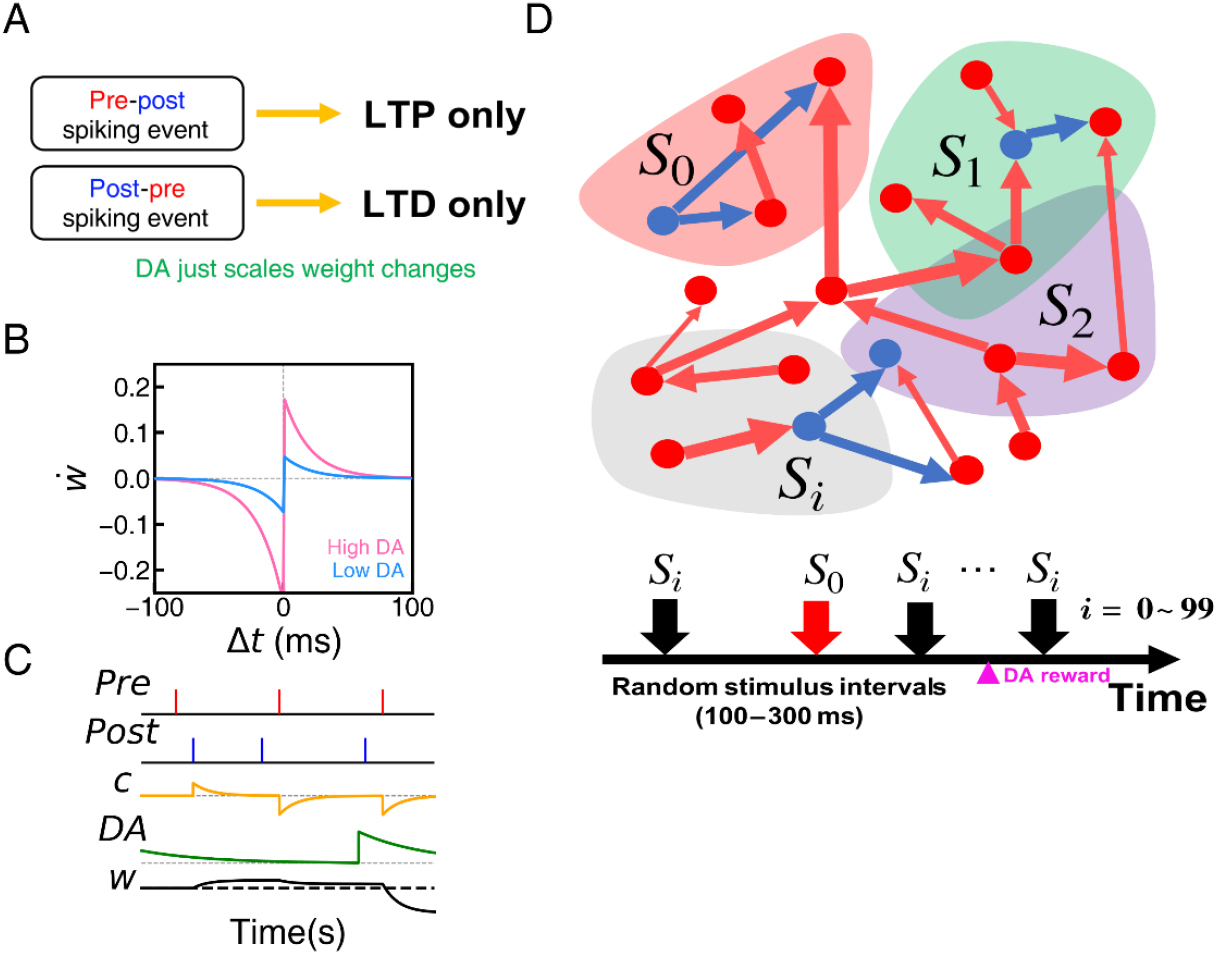
Schematic illustration of the original Pavlovian conditioning paradigm in a spiking neural network. (A) Pre-post spike pair events induce LTP, while post-pre spike pair events induce LTD. STDP function curves under low (light blue) and high (pink) DA concentrations. (C) Exemplary time course of pre- and post spiking events (Pre & Post), eligibility trace (*c*), DA concentration, and synaptic weight (*w*). (D) Model spiking neural network composed of excitatory (red) and inhibitory (blue) neurons organized into slightly overlapping subgroups (*S*_*i*_), stimulated in a random sequence with variable interstimulus intervals. Only the stimulus delivered to subgroup *S*_0_ is paired with distal dopamine reward (filled, violet triangle).

Such nonlinear modulation of synaptic plasticity is thought to contribute to the widely observed inverted-U relationship between DA concentration and learning performance, whereby both insufficient and excessive DA impair learning and cognitive function (Cools & D’Esposito, 2011; Westbrook, Booij, Cools, & Narayanan, 2022). Motivated by these findings, we propose a DA-modulated STDP model in which increasing DA progressively biases synaptic modification toward potentiation, while receptor saturation limits further DA effects beyond a critical concentration. We demonstrate that these two biologically motivated mechanisms define a finite, biologically realistic DA window for Pavlovian conditioning in recurrent spiking neural networks.

## 2 Results

### 2.1 Original DA-modulated STDP rule and its learning performance on DA level

We first evaluated the learning performance of the original Izhikevich DA-modulated STDP model in a recurrent spiking neural network consisting of 2,000 excitatory and inhibitory neurons performing reward-based conditioning with spatially patterned stimuli. To isolate the effect of dopamine, the initial network configuration and the sequence of presented stimuli were kept identical across all simulations. Representative network responses at three reward DA concentrations are shown in Fig. 2A–C.

**Fig. 2.**
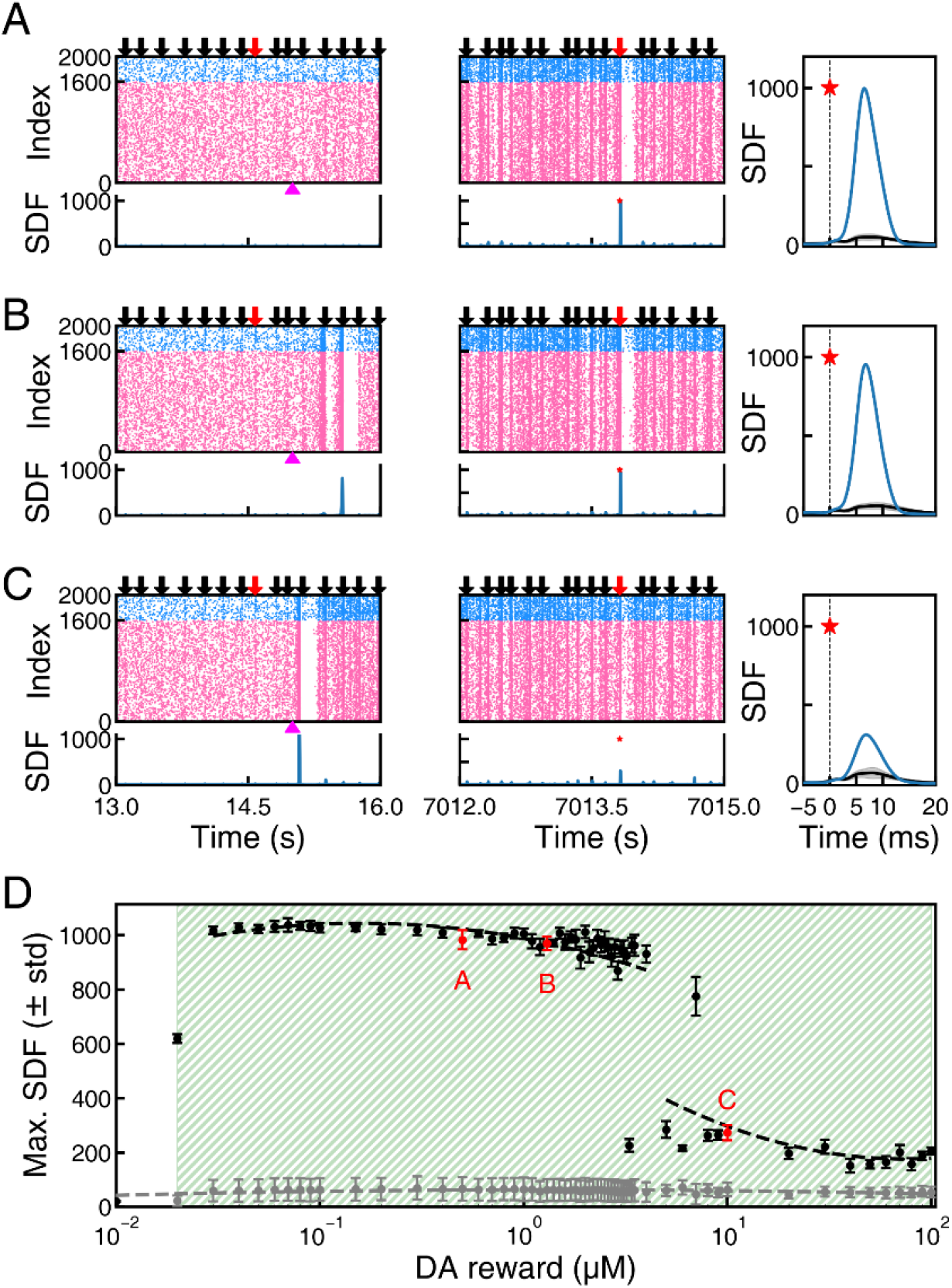
Assessing learning performance of the original Izhikevich DA-modulated STDP model across reward DA concentrations. Spike raster plots of excitatory (pink) and inhibitory (blue) neurons during the early (left column) and late (middle column) phases of conditioning at reward DA concentrations of (A) 0.50 *µ*M, (B) 1.30 *µ*M, and (C) 10.00 *µ*M. Corresponding spike-density functions (SDFs) are shown beneath each raster plot. The right column shows enlarged SDFs during the post-conditioning test phase for reward-associated stimuli (blue) and the mean response to non-reward-associated stimuli (black; gray shading indicates the standard deviation). The star denotes the time of stimulus presentation. (D) Peak SDF as a function of reward DA concentration for reward-associated (black) and non-reward-associated (gray) stimuli during the test phase. Dashed curves are spline fits provided only as visual guides. Error bars were calculated from 11 reward-associated trials and 939 non-reward-associated trials. Red labels (A–C) indicate the DA concentrations illustrated in panels (A–C). The green hatched region denotes the range of reward DA concentrations yielding an AUC≥ 0.9. All simulations used the same initial network configuration and identical stimulus presentation sequence.

At a moderate reward DA concentration ([DA] = 0.50 *µ*M; Fig. 2A), responses to reward-associated (red arrow) and non-reward-associated (black arrows) stimuli were indistinguishable during the early stage of conditioning. In all cases, stimulus-evoked activity remained confined to the directly stimulated neuronal population, resulting in similarly small spike-density functions (SDFs). Following conditioning, however, reward-associated stimuli recruited nearly the entire network, generating synchronized burst responses with substantially larger SDF peaks, whereas non-reward-associated stimuli continued to activate only a small subset of neurons.

Remarkably, increasing the reward DA concentration to 1.30 *µ*M produced nearly identical conditioning results (Fig. 2B). Although occasional nonspecific burst responses appeared during the early phase of conditioning, the fully conditioned network remained highly selective, exhibiting robust burst responses only to reward-associated stimuli.

Even at an exceptionally high DA concentration ([DA] = 10.00 *µ*M; Fig. 2C), the conditioned network retained strong stimulus selectivity. Although the burst response to the reward-associated stimulus was reduced compared with the previous cases, it remained readily distinguishable from responses to non-reward-associated stimuli, with the area under the receiver operating characteristic curve (AUC) exceeding 0.9. Across the entire range of tested reward DA concentrations (0.02–100 *µ*M) explored, the original model consistently achieved successful conditioning (Fig. 2D), maintaining an AUC above 0.9. Notably, however, the peak SDF exhibited an abrupt decrease at DA concentrations of approximately 7–8 *µ*M, separating the data into two distinct branches (black symbols, indicated by the dashed lines).

### 2.2 Modified DA-modulated STDP rule

Our model is based on the hypothesis that dopamine not only gates STDP but also nonlinearly reshapes the STDP window itself (Fig. 3A), consistent with experimental observations (Brzosko et al., 2015; Zhang et al., 2009). We hypothesize that this dopamine-dependent deformation, together with receptor saturation, restricts reward-based learning to a biologically realistic range of dopamine concentrations. Specifically, the model progressively biases the STDP window toward potentiation as dopamine levels increase (Fig. 3B), while receptor saturation limits further deformation beyond a critical dopamine concentration. To implement this mechanism, the eligibility trace is decomposed into pre–post and post–pre spike interaction components. At low dopamine concentrations, pre–post and post–pre spike pairs induce LTP and LTD, respectively. As dopamine levels increase, however, the depression associated with post–pre spike pairs is progressively weakened and eventually converted into LTP (Fig. 3C). To achieve this continuous transition, we assume that the rate of change of the synaptic weight is proportional to a quadratic function of DA (see Methods for details).

**Fig. 3.**
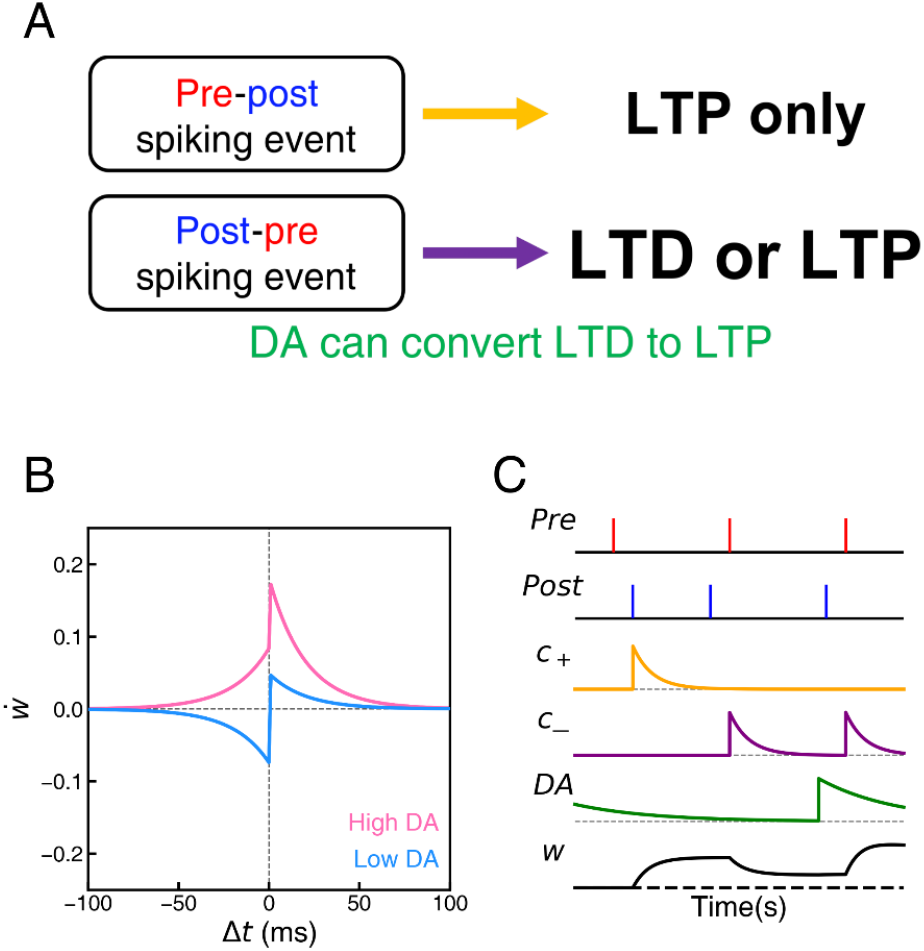
Modified dopamine-modulated STDP model. (A,B) Schematic illustration of the proposed dopamine-dependent deformation of the STDP window. At low DA concentrations, as in the original model, post–pre spike pairings induce long-term depression (LTD), whereas pre–post spike pairings induce long-term potentiation (LTP). As DA concentration increases, the depression associated with post–pre spike pairings is progressively weakened and eventually converted into LTP, thereby reshaping the STDP window rather than simply scaling its magnitude. (C) Representative time courses of the model variables governed by the modified plasticity rule, including the eligibility traces for pre–post (*c*_+_) and post–pre (*c*_−_) spike interactions, dopamine concentration (DA), and synaptic weight (*w*) in response to pre- and postsynaptic spike events.

### 2.3 Learning performance of the modified DA-modulated STDP model across dopamine concentrations

The original Izhikevich model supported successful conditioning over an unrealistically broad range of reward DA concentrations (Fig. 2), failing to reproduce the experimentally observed inverted-U relationship between dopamine and learning. We therefore examined whether the proposed dopamine-dependent deformation of the STDP window, together with receptor saturation, restricts successful conditioning to a biologically realistic DA range. Figure 4 summarizes the results obtained with the modified learning rule using the same initial random network configuration and identical stimulus presentation sequence employed in Fig. 2.

**Fig. 4.**
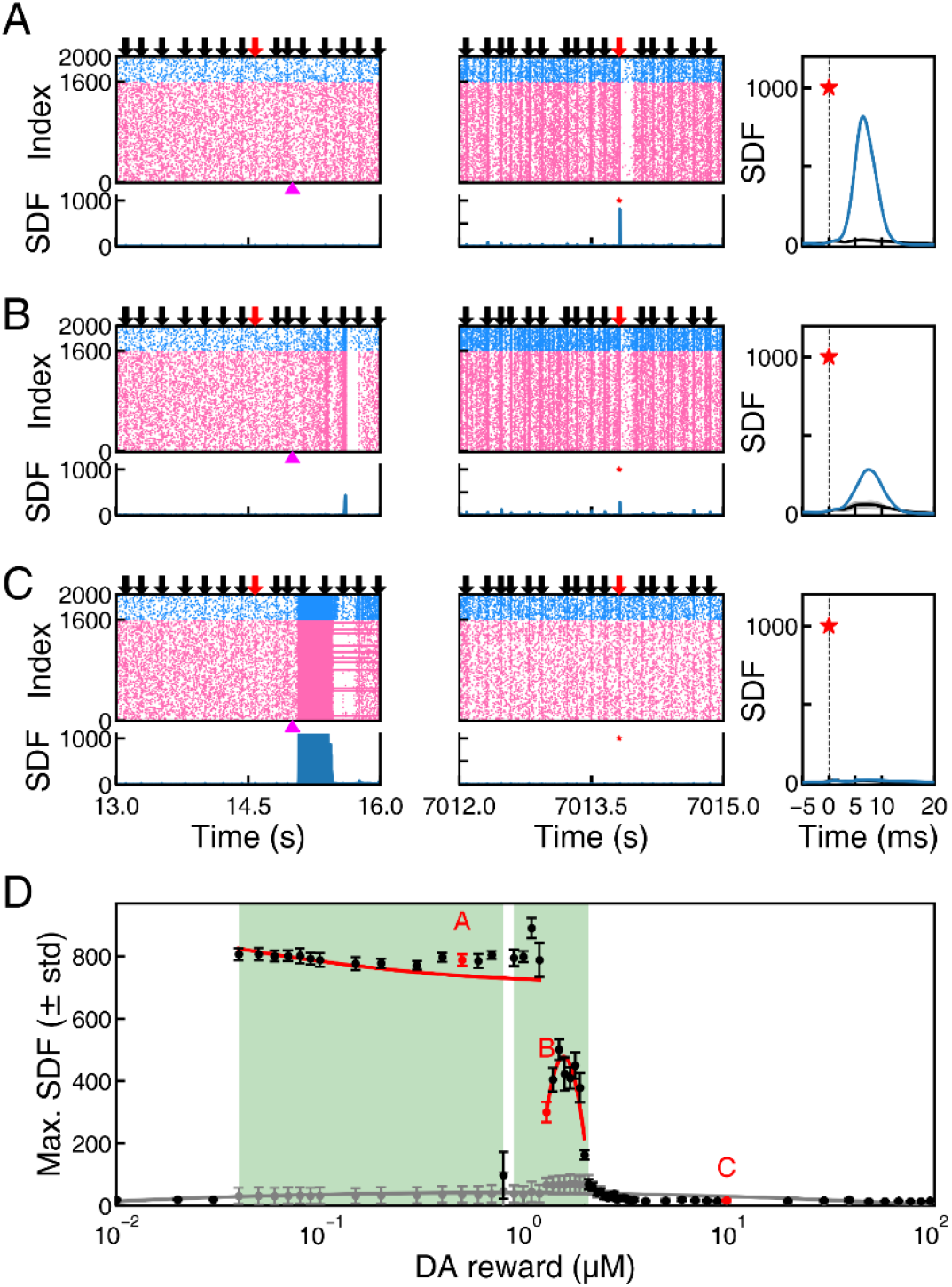
Learning performance of the modified DA-modulated STDP model across reward DA concentrations. The layout and plotting conventions are identical to those of Fig. 2. Results are shown for reward DA concentrations of (A) 0.50 *µ*M, (B) 1.30 *µ*M, and (C) 10.00 *µ*M. (D) Peak spike-density-function (SDF) values during the post-conditioning test phase as a function of reward DA concentration. Red labels indicate the conditions illustrated in panels (A)–(C). The green hatched region denotes the range of reward DA concentrations yielding an AUC ≥ 0.9. All simulations used the same initial network configuration and identical stimulus presentation sequence as in Fig. 2.

At a biologically relevant reward DA concentration ([DA] = 0.50 *µ*M; Fig. 4A), the modified model exhibited learning behavior similar to that of the original model. During the early phase of conditioning, both reward-associated and non-reward-associated stimuli elicited activity largely confined to the directly stimulated neuronal population, producing small and nearly identical spike-density-function (SDF) responses. Following conditioning, however, reward-associated stimuli recruited most of the network, generating synchronized burst responses with substantially larger SDF peaks, whereas non-reward-associated stimuli continued to evoke only weak local activity.

When a higher reward DA concentration ([DA] = 1.30 *µ*M) was used, the early phase of conditioning closely resembled that of the original Izhikevich model, with no appreciable difference in network responses to reward-associated and non-reward-associated stimuli (Fig. 4B, left column). In contrast, the post-conditioning test phase exhibited markedly different behavior. Reward-associated stimuli elicited substantially weaker burst responses than in the original model at the same DA concentration. Instead, the responses closely resembled those produced by the original Izhikevich model only under an unrealistically high reward DA concentration ([DA] = 10.00 *µ*M; Fig. 2C).

When a reward DA concentration of [DA] = 10.00 *µ*M was used, the modified model exhibited markedly different behavior from the original Izhikevich model throughout both the training and test phases (Fig. 4C). During the early phase of conditioning, reward-associated and non-reward-associated stimuli again evoked spiking largely confined to the directly stimulated neuronal population. Following each DA reward delivery, however, the network entered a prolonged period of high-frequency population activity, with most neurons remaining active for an extended duration. This behavior contrasts with that of the original model, in which DA reward delivery elicited only brief, synchronized bursts during early training (Fig. 2C). After conditioning, reward-associated and non-reward-associated stimuli both evoked only weak, locally confined responses, yielding virtually indistinguishable SDF profiles. Thus, conditioning failed completely at this high DA concentration.

### 2.4 Evolution of Network Structure

In our previous study (Jeong & Lee, 2025), we showed that a scatter plot of the total incoming synaptic weight ( Σ *W*_in_) versus the total outgoing synaptic weight ( Σ *W*_out_) for all excitatory neurons provides a useful visualization of network structural evolution during conditioning. We therefore used this analysis to compare network developmental trajectories across different reward DA concentrations. At a moderate reward DA concentration ([DA] = 0.50 *µ*M), the network evolved similarly to the original Izhikevich model (Fig. 5A). By *t* = 1,200 s, a strong negative correlation had developed between Σ *W*_out_ and Σ *W*_in_, indicating the emergence of a pronounced feedforward backbone. Within this organization, activity evoked by the reward-associated stimulus (*S*_0_) propagates along the backbone from neurons with large outgoing and small incoming synaptic weights (upper left) toward neurons with small outgoing and large incoming weights (lower right). Neurons belonging to the *S*_0_ subpopulation are high-lighted in red in Fig. 5. The emergence of this negative correlation, together with the tight clustering of the *S*_0_ neurons, constitutes a hallmark of successful conditioning. This network organization formed rapidly during the early phase of conditioning and remained stable throughout the remainder of the simulation (*t* = 3,600 s).

**Fig. 5.**
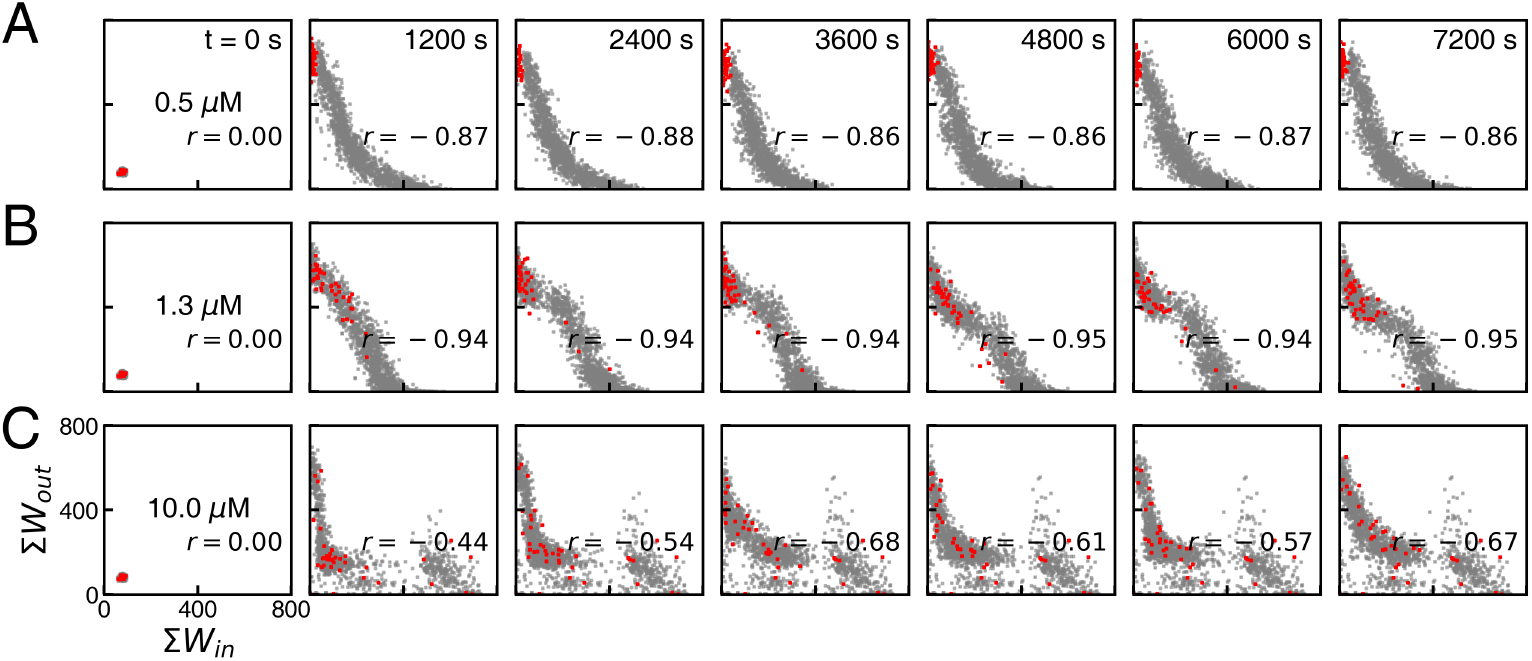
Temporal evolution of network structure under the modified DA-modulated STDP rule. Network development is visualized in the Σ *W*_out_ versus Σ *W*_in_ plane. The initial network configuration and stimulus presentation sequence were identical to those used in Fig. 4. Each point represents an excitatory neuron. Neurons belonging to the reward-associated (*S*_0_) population are shown in red, whereas all other neurons are shown as semi-transparent gray points. Panels (A)– correspond to reward DA concentrations of 0.50 *µ*M, 1.30 *µ*M, and 10.00 *µ*M, respectively. As the reward DA concentration increases, the strong negative correlation between Σ *W*_out_ and Σ *W*_in_ gradually weakens, and the *S*_0_ neurons become progressively less clustered. The Spearman correlation coefficient *r* measures the strength and direction of the monotonic relationship between two variables by computing the Pearson correlation coefficient on their rank-transformed values.

At a higher reward DA concentration ([*DA*] = 1.30 *µ*M; Fig. 5B), the network followed a noticeably different developmental trajectory. As in the 0.50 *µ*M case, a pronounced feedforward backbone again emerged, with the *S*_0_ neurons clustering in the upper-left region of the scatter plot. Compared to the lower-DA case ([*DA*] = 0.50 *µ*M), however, the *S*_0_ cluster was less compact and fluctuated significantly over time. The negative Spearman correlation *r* between *W*_out_ and *W*_in_ remained fairly steady beyond *t* = 1,200 s, at approximately *r* = − 0.95 — notably larger in magnitude than the *r* = − 0.87 observed for [*DA*] = 0.50 *µ*M. *r* is the value of Spearman correlation.

At a very high reward DA concentration ([*DA*] = 10.00 *µ*M; Fig. 5C), the network largely failed to develop a coherent feedforward backbone. Instead, neurons dispersed into multiple clusters across the two-dimensional scatter plot. More importantly, the *S*_0_ neurons receiving reward-associated stimuli no longer formed a stable cluster in the upper-left region but instead migrated irregularly throughout conditioning and became broadly dispersed. This behavior contrasts sharply with that of the original Izhikevich model, in which a well-defined feedforward backbone emerged consistently across the entire range of tested DA concentrations, and the *S*_0_ neurons remained tightly clustered in the upper-left region of the scatter plot (Supplementary Fig. S1). Notably, the network was also unstable throughout the conditioning process: both the degree of *S*_0_ subpopulation dispersion and the Spearman correlation *r* between Σ *W*_out_ and Σ *W*_in_ fluctuated substantially.

### 2.5 Comparison of modified and original models across dopamine concentrations

While Fig. 4D illustrates learning performance from a single representative simulation, Fig. 6 summarizes the results across *N* = 11 independent simulations with different initial network configurations and stimulus presentation sequences. For each simulation, the mean and standard deviation of the SDF peak during the post-conditioning test phase were first calculated across repeated stimulus presentations. The per-simulation means were then averaged across all 11 simulations and are presented as mean *±* standard error of the mean (SEM). The inset in Fig. 6B shows the individual simulation results (mean *±* standard deviation) within the bistability-like regime without averaging across simulations.

**Fig. 6.**
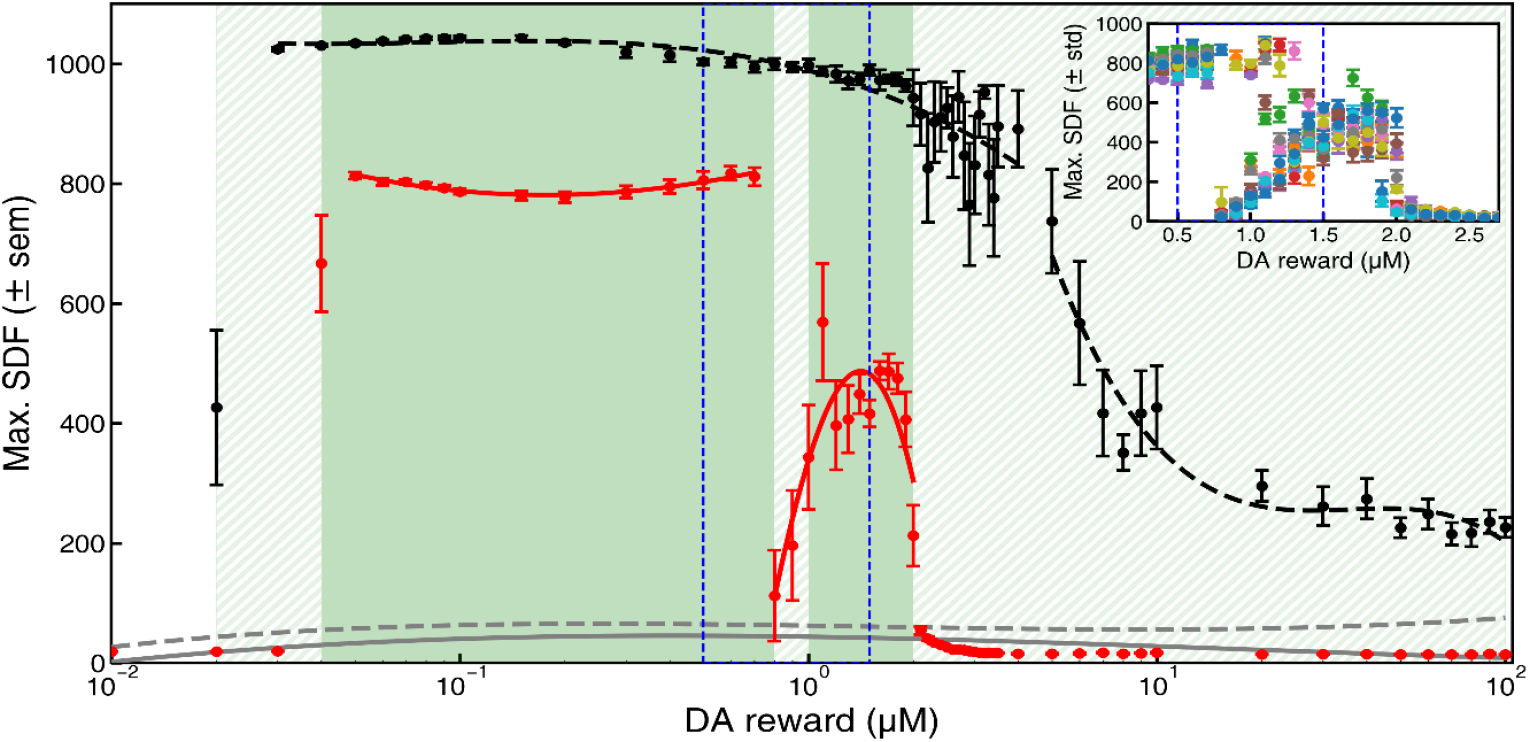
Comparison of learning performance between the original and modified DA-modulated STDP models. (A) Mean spike-density-function (SDF) peak during the post-conditioning test phase as a function of reward DA concentration for the original Izhikevich model (black circles and dashed line) and the modified model (red circles and solid line). Responses to non-reward-associated stimuli are shown in gray. Green shaded regions indicate DA concentrations yielding an AUC≥ 0.9 for each model. (B) Expanded view of the bistability-like regime in the modified model, bounded by the vertical blue dashed lines.

The modified model produced robust learning over a biologically realistic DA range (0.04–0.70 *µ*M), exhibiting consistently large SDF peak differences between reward-associated and non-reward-associated stimuli. At 0.80 *µ*M, learning performance declined sharply, followed by a partial recovery and a subsequent decrease up to 2.00 *µ*M, beyond which responses to reward-associated and non-reward-associated stimuli became indistinguishable. In contrast, the original Izhikevich model maintained strong learning performance over a much broader DA range (0.03–4.00 *µ*M), followed by a gradual decline up to 10.00 *µ*M while still preserving stimulus selectivity. The non-monotonic behavior observed between 0.80 and 2.00 *µ*M varied among simulations and was therefore examined in greater detail (Fig. 7).

**Fig. 7.**
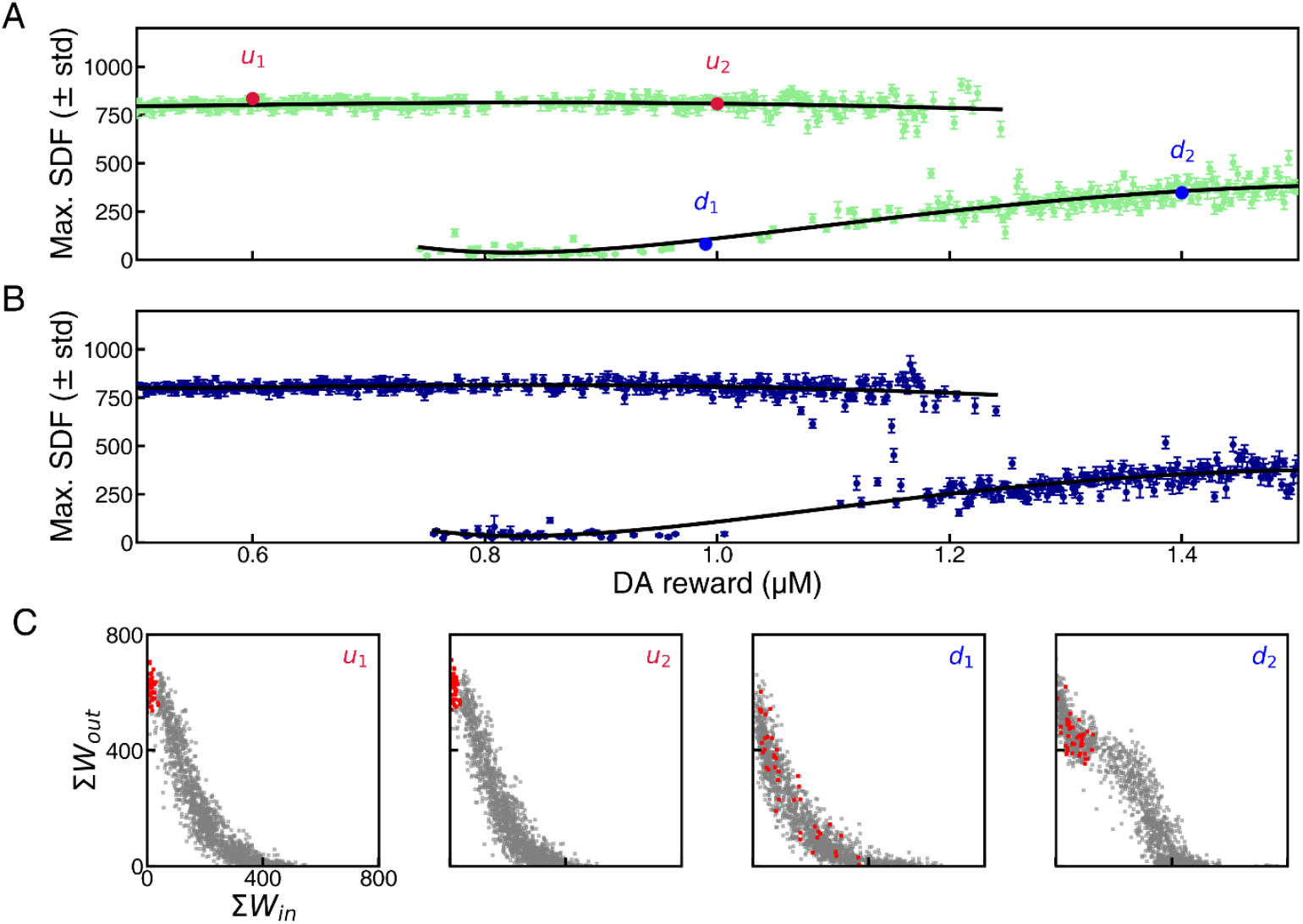
Multistability near the transition between successful and unsuccessful conditioning. (A) SDF peak as a function of reward DA concentration during the ascending sweep. Representative upstate (*u*_1_, *u*_2_; red) and downstate (*d*_1_, *d*_2_; blue) network states are highlighted. Black curves denote piecewise spline fits to the upper and lower branches. (B) Corresponding results for the descending sweep. (C) Network structures corresponding to the highlighted states in (A), visualized in the Σ*W*_out_ versus Σ*W*_in_ plane using the same conventions as in Fig. 5.

### 2.6 Multi-stability in the transition zone of DA

To determine whether the complex behavior of peak SDF values observed between 0.80 and 2.00 *µ*M reflects genuine multistability associated with the conditioning, we performed a sequential continuation analysis. Instead of initializing each simulation with a new random network, the conditioned network obtained at one reward DA concentration was used as the initial condition for the immediate next simulation: Reward DA concentration was then varied in small increments from 0.50 to 1.50 *µ*M and subsequently decreased back to 0.50 *µ*M, while recording the SDF peak after each conditioning run (Fig. 7A,B). The black curves represent piecewise spline fits separately applied to the upper and lower branches using a fixed SDF threshold of 550.

Both the ascending and descending sweeps exhibited abrupt transitions between two discrete network states, with no evidence of a continuous intermediate state. Representative points from the upper and lower branches (*u*_1_, *u*_2_, *d*_1_, and *d*_2_) were selected for structural analysis in Fig. 7C.

Across both sweeps, the network occupied either an upstate or a downstate. The upstate persisted from 0.50 *µ*M to approximately 1.20 *µ*M. In this state, the network developed a stable feedforward backbone, and neurons receiving reward-associated stimuli remained clustered in the upper-left region of the weight space. The downstate emerged from approximately 0.80 to 1.50 *µ*M. At the lower end of this range, the network exhibited a compressed feedforward structure shifted toward the lower-left region, with reward-associated neurons broadly dispersed. As reward DA concentration increased, these neurons gradually migrated toward the upper-left region, whereas intermediate neurons within the feedforward hierarchy shifted toward the upper-right direction, producing an increasingly unstable feedforward organization.

## 3 Discussion

The original Izhikevich model provides an elegant framework for reward-based learning, in which spatiotemporal neuronal activity is associated with temporally delayed DA reward through an eligibility trace mechanism. However, because DA acts only as a multiplicative factor that uniformly scales the STDP window, the balance between long-term potentiation (LTP) and long-term depression (LTD) remains unchanged over the entire range of DA concentrations. Consequently, the model supports successful conditioning even under biologically unrealistic DA levels. To overcome this limitation and provide a more biologically grounded framework for modeling dopaminergic learning and neurological disorders, we proposed a modified DA-modulated STDP rule in which DA progressively reshapes the STDP window while receptor saturation limits its effect at high concentrations.

Both the original and modified models fail to produce conditioning when the reward DA concentration is too low, because DA is required to reinforce both potentiation and depression through the eligibility trace mechanism. At the opposite extreme, however, the two models behave fundamentally differently. In the original model, increasing DA simply amplifies synaptic modification while preserving the relative balance between LTD and LTP. Consequently, the feedforward organization established by the rewarded *S*_0_ stimulus remains robust over an unrealistically broad DA range. In the modified model, by contrast, elevated DA progressively suppresses LTD until the STDP window becomes dominated by potentiation regardless of the relative timing of pre- and postsynaptic spikes. As a result, spike-timing-dependent causality—the mechanism by which rewarded *S*_0_ activity is preferentially reinforced—is gradually lost. Once this temporal selectivity disappears, the rewarded input no longer provides a privileged direction for network self-organization. Instead, network structure appears to emerge primarily through internally generated dynamics, producing the fragmented clustering observed in Fig. 5C. Interestingly, this organization resembles the clustered structures previously observed in recurrent networks that self-organize into superburst-generating states (Kim & Lee, 2022), suggesting that excessive DA may shift the dominant organizing principle from externally driven learning to intrinsic network dynamics.

Another intriguing finding of the modified model is the emergence of a narrow transition regime exhibiting multistability. Within this regime, the network converged to one of two distinct structural organizations despite identical reward DA concentrations. Nevertheless, the observed behavior does not fully satisfy the classical notion of bistability in low-dimensional dynamical systems. Rather than remaining in one attractor until a critical parameter threshold is crossed, the network occasionally switched between the two states following only small changes in reward DA concentration, even well within the transition region. We speculate that this sensitivity reflects the high dimensionality of the recurrent network, in which the competing attractors possess complex, intertwined basins of attraction. Under such conditions, even a small change in DA concentration may alter the trajectory of the conditioning process sufficiently to direct the network toward a different attractor. It is important to emphasize, however, that each change in DA concentration was accompanied by an additional conditioning period under the new parameter value. Therefore, the observed transitions should not be interpreted as the immediate consequence of an infinitesimal parameter perturbation, but rather as the outcome of continued synaptic reorganization under a slightly modified DA environment.

Our modified DA-modulated STDP model is fundamentally a phenomenological extension of the original model proposed by Izhikevich (2007). Rather than explicitly modeling the underlying molecular signaling pathways, it qualitatively reproduces the experimentally observed dopamine-dependent deformation of the STDP window reported by Zhang et al. (2009) and Brzosko et al. (2015). To capture these observations, we introduced two eligibility traces, replacing the single eligibility trace used in the original model. To capture these experimental observations, we introduced two eligibility traces in place of the single trace used in the original model. Although purely phenomenological, these two variables are intended to represent the functionally distinct intracellular processes leading to LTP and LTD, both of which are critically regulated by intracellular Ca^2^+ signaling. (Graupner & Brunel, 2012; Lisman, 2001; Shouval, Bear, & Cooper, 2002) The associated model parameters were chosen to reproduce the experimentally observed dependence of the STDP window on dopamine concentration, rather than to represent specific biochemical reaction rates.

It should also be emphasized that DA-modulated STDP is unlikely to represent the sole mechanism underlying dopamine-dependent learning. Synaptic plasticity depends not only on spike timing but also on numerous other factors, including firing rate, post-synaptic membrane potential, intracellular ionic dynamics, and the biochemical state of the synapse (Citri & Malenka, 2008; Feldman, 2012). Moreover, dopamine generally does not act in isolation but interacts with other neuromodulators, particularly acetylcholine (ACh), to regulate corticostriatal synaptic plasticity and learning. (Calabresi et al., 2007; Kreitzer & Malenka, 2008) Incorporating these additional mechanisms will be essential for developing more biologically realistic models of reward-based learning and its dysfunction in neurological disorders.

## 4 Methods

### 4.1 Network architecture

The recurrent spiking neural network consisted of 2,000 neurons, comprising 1,600 excitatory and 400 inhibitory neurons that are randomly connected with a sparsity of 0.1. Each neuron was modeled using the Izhikevich neuron model, governed by the following differential equations:

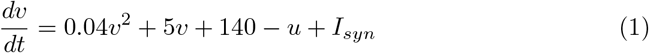

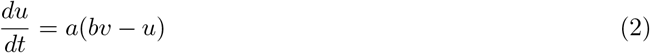

where *v* denotes the membrane potential, *u* is a membrane recovery variable, and *I*_*syn*_ is the synaptic input current. A spike is generated when *v* ≥ 30 mV, after which the variables are reset according to:

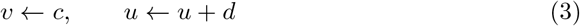

Excitatory and inhibitory neurons were parameterized as (*a, b, c, d*) = (0.02, 0.2, −65, 8) and (0.1, 0.2, −65, 2), respectively, with all neurons initialized at *v* =− 65 mV and *u* = − 13. One hundred stimulus subgroups (*S*_0_ through *S*_99_) were defined by randomly assigning 50 neurons (40 excitatory and 10 inhibitory) to each subgroup, with overlaps between subgroups permitted. All neurons were connected with a fixed connection probability of *p* = 0.1. Excitatory synaptic weights were initialized to *w*_0_ = 0.5 and bounded within [0, 4.0]. Inhibitory synaptic weights were fixed at *w*_*I*_ = −0.5 throughout the simulation.

### 4.2 DA-modulated STDP rule

Dopamine concentration *DA* at each neuron was modeled as an exponentially decaying variable that increases upon reward delivery:

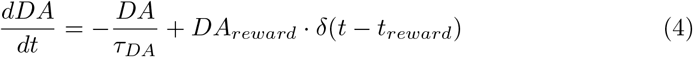

where *τ*_*DA*_ = 200 ms is the dopamine decay time constant and *DA*_*reward*_ is the reward dopamine concentration, which was held constant throughout the training phase.

In the original Izhikevich model, synaptic plasticity is mediated by a single eligibility trace *c*, which tracks the history of pre- and postsynaptic spike interactions:

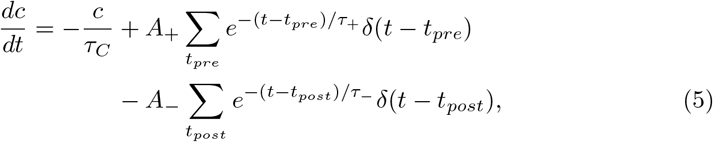

where *τ*_*C*_ = 1000 ms is the eligibility trace decay time constant, *τ*_+_ = *τ*_−_ = 20 ms are the STDP time constants, and *A*_+_ = 0.1, *A*_−_ = 0.15 are the potentiation and depression amplitudes. Synaptic weight updates are then governed by:

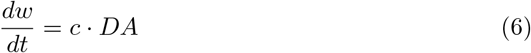

In this formulation, dopamine acts as a simple multiplicative gate, scaling the rate of synaptic modification linearly with dopamine concentration without altering the shape or polarity of the STDP window.

In the modified model, the eligibility trace is separated into two independent components, *c*_+_ and *c*_−_, tracking pre-post and post-pre spike interactions respectively:

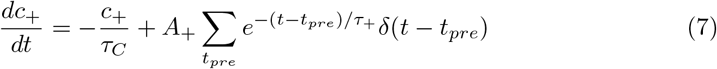

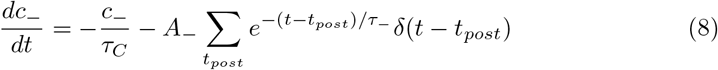

Dopamine nonlinearly deforms the STDP window through a saturating transformation of dopamine concentration:

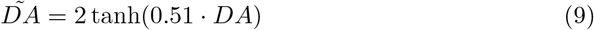

This transformation implements receptor saturation, ensuring that the influence of dopamine on synaptic plasticity does not increase indefinitely at high concentrations. A key change from the previous model is that synaptic weight updates are now assumed to be governed by:

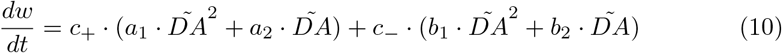

where *a*_1_, *a*_2_, *b*_1_, *b*_2_ are parameters controlling the dopamine-dependent deformation of the STDP window. Parameters were set to *a*_1_ = 0, *a*_2_ = 1, *b*_1_ = 1, *b*_2_ = − 1.5, such that pre-post spike interactions consistently induce LTP as in the original model, while post-pre spike interactions produce LTD at low dopamine concentrations but transition toward LTP as dopamine concentration increases beyond a critical value (of *DA* = 1.5 for the chosen values of *b*_1_ = 1, *b*_2_ = −1.5). To our knowledge, this is the first model to describe such a transition.

### 4.3 Stimulation and reward protocol

Simulations were run for 7,200 seconds with a temporal resolution of 1 ms. At each timestep, all neurons received a background noise current drawn from a uniform distribution over [− 6.5, +6.5].

During the training phase (0 – 7,000 s), stimuli were delivered at random intervals uniformly distributed between 100 and 300 ms. At each stimulus event, a randomly selected subgroup received an input current of 40 for a single timestep. When the selected subgroup was *S*_0_, a dopamine reward of concentration *DA*_*reward*_ was delivered at a random delay uniformly distributed between 0 and 1,000 ms following stimulus onset. To ensure that a reward-associated stimulus (*S*_0_) was present in both the early and late recording windows, a single *S*_0_ presentation was inserted at fixed positions within each recording period. During the post-training test phase (7,000 – 7,200 s), stimuli were delivered under the same protocol but dopamine reward was withheld. Network spiking responses during this phase were used to evaluate learning performance, including SDF peak values and stimulus discriminability.

### 4.4 Data analysis

Network spiking activity was quantified using the spike density function (SDF), computed by binning spike times into a histogram with a bin width of 1 ms and smoothing with a Gaussian kernel of standard deviation *σ* = 1 ms:

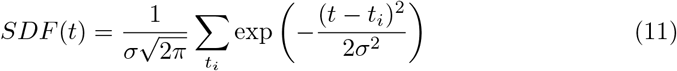

Learning performance was assessed by comparing SDF peak values evoked by reward-associated and non-reward-associated stimuli during the post-training test phase. For each stimulus presentation, the maximum SDF value within a 10 ms window following stimulus onset was extracted as the SDF peak value.

Stimulus discriminability was quantified using the area under the receiver operating characteristic curve (AUC), computed nonparametrically via the Mann-Whitney *U* statistic, following the ROC-based discriminability framework established in neurophysiology (Britten, Shadlen, Newsome, & Movshon, 1992):

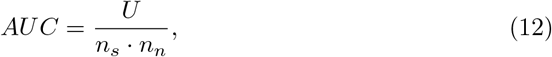

where *U* is the Mann-Whitney *U* statistic comparing SDF peak distributions of reward-associated stimulus *S*_0_ (*n*_*s*_ trials) against non-reward-associated stimuli (*n*_*n*_ trials), with the alternative hypothesis that *S*_0_ responses are greater. An AUC ≥ 0.9 was defined as selective, indicating clear discrimination between reward-associated and non-reward-associated stimuli (Mandrekar, 2010). For the sequential simulation analysis, network states were classified into upper and lower states using a fixed SDF peak threshold of 550.

### 4.5 Simulation environment

All simulations were implemented using PyGeNN (version 4.9.0), a Python interface to the GPU-enhanced Neuronal Networks (GeNN) framework, which enables efficient large-scale spiking neural network simulations on GPU hardware. Simulations were executed on a system equipped with two NVIDIA GeForce RTX 3090 GPUs. Simulation code is available at https://doi.org/10.5281/zenodo.20443391.

## Supplementary information

Supplementary Figure S1 accompanies this article.

**Fig. S1.**
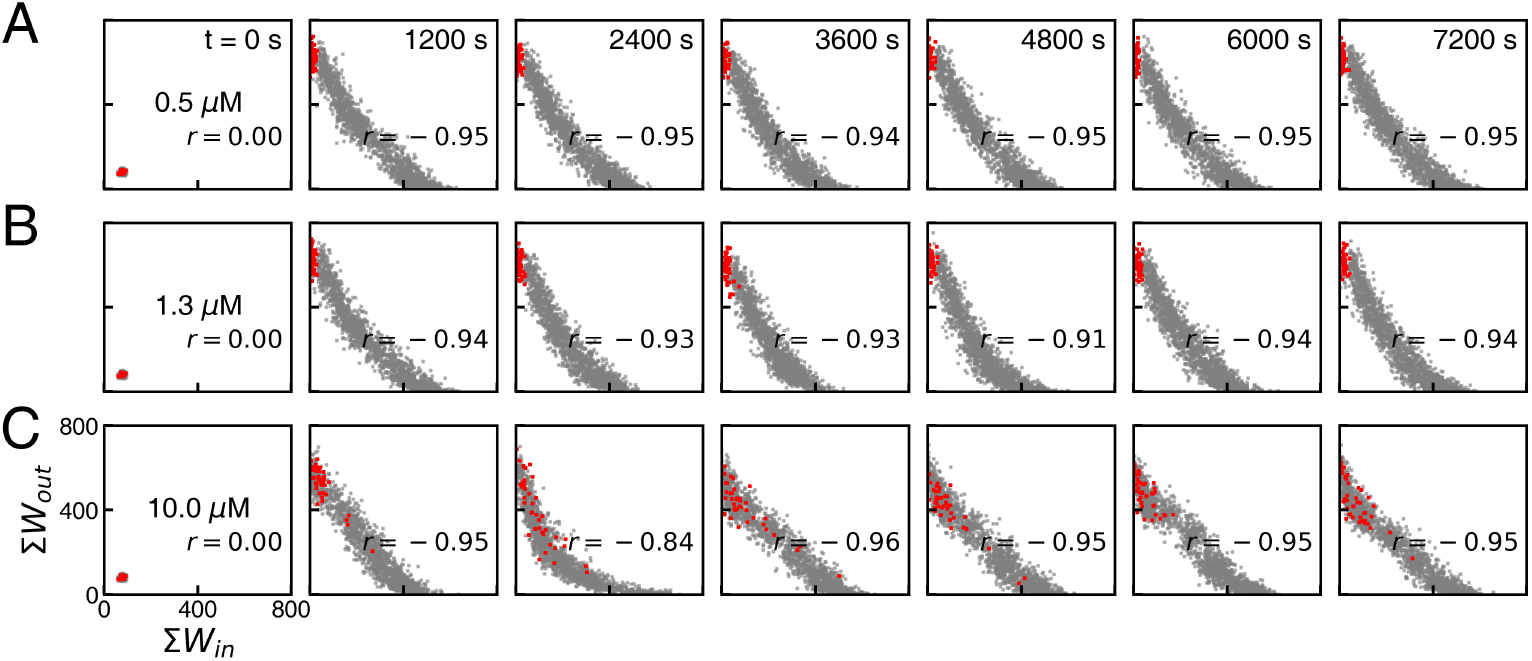
Temporal evolution of network organization during the Izhikevich conditioning protocol based on the conventional three-factor learning rule, visualized in Σ*W*_in_ vs. Σ*W*_out_ space. The network composition and stimulus sequence are identical to those used in Fig. 4. All visualization conventions are the same as in Fig. 5. Reward dopamine concentrations were set to (A) 0.50 *µ*M, (B) 1.30 *µ*M, and (C) 10.00 *µ*M. Note that even for DA = 10.00 *µ*M, a negative correlation between weight-in and weight-out is evident and red points remain prominently aggregated in the upper-left corner.

## Declarations

### Ethics approval

Not applicable. This study did not involve human participants or animal subjects; all data were generated through computational simulation.

### Data availability

The simulation code supporting the findings of this study is openly available on Zenodo at https://doi.org/10.5281/zenodo.20443391 (mirrored from GitHub: koruniv1098-lgtm/modified-dopamine-stdp-pavlovian).

### Author contributions

KJL conceived the project and advised WP to perform numerical simulations and carry data analysis. WP and KJL wrote the paper together.

