## Supplementary figures and images for "Biologically Plausible Dopamine-Modulated STDP Model of Pavlovian Learning in Spiking Neural Networks"

### Supplementa Figure 1

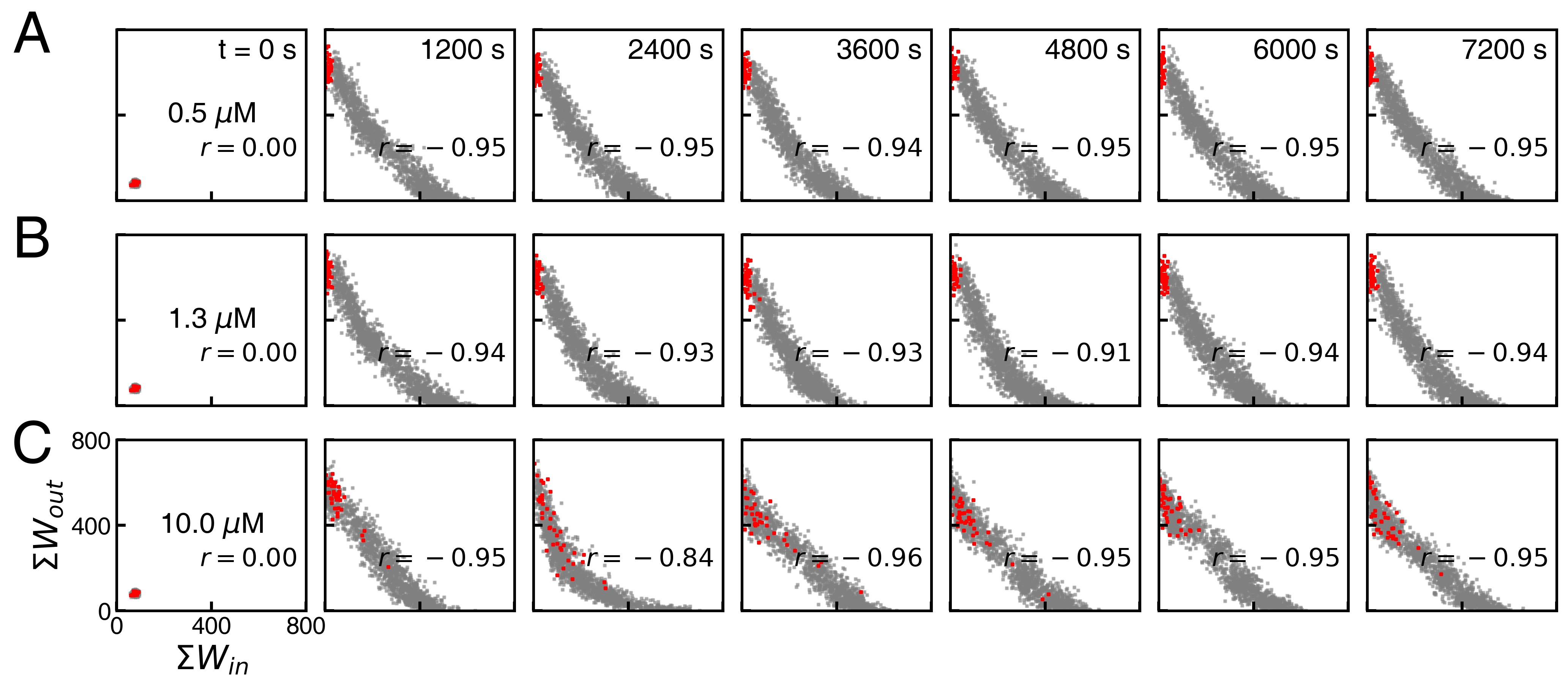
